# Mid-infrared light nonthermally produces anti-aging effects through cellular efficiency enhancement in living organisms

**DOI:** 10.64898/2026.02.08.704722

**Authors:** Changsheng Shao, Daoling Peng, Yunlong Zhao, Yousheng Shu, Songlin Zhuang, Qing Huang, Bo Song

## Abstract

Aging is commonly viewed as a consequence of cumulative molecular damage and declining stress resilience. However, this perspective overlooks the possibility that aging may primarily arise from an efficiency loss of integrated cellular operation. Here, we show that aging can be systemically prevented by enhancing cellular efficiency using frequency-specific mid-infrared (MIR) light, as revealed by organismal-, cellular- and molecular-level analyses of *Caenorhabditis* elegans. Remarkably, super-weak 34-THz MIR light (∼1 μW mm^−^²) prolonged the median lifespan of worms by 60%, delaying aging onset and preventing abrupt “cliff-edge” mortality, without detectable thermal effects. At the cellular level, MIR exposure enhanced global gene transcription in parallel with the establishment of a mitochondrial state of high-efficiency energy metabolism, together preserving youthful cellular homoeostasis during aging. These cellular effects were associated with vibrational modes of phosphate groups in nucleic acids and mitochondrial phospholipids within the 33–35 THz range, providing a frequency-matched molecular context for MIR modulation. Together, our results support an efficiency-first, systemic anti-aging model in which frequency-specific MIR light enhances the integrated kinetic efficiency of cellular systems, spanning gene transcriptional dynamics and mitochondrial bioenergetic flux, rather than the activation of damage-repair or stress-response pathways. These findings advance our understanding of light-matter interactions in living systems and establish a non-biochemical, physical strategy for modulating aging.

---

Population aging is rapidly reshaping global health, as age-related functional decline underlies most chronic diseases and limits healthspan worldwide (1). Understanding and modulating the biological basis of aging has therefore become a central challenge across medicine, biology and society. Aging has long been interpreted through a framework of cumulative molecular damage and compensatory stress responses (2). Within this view, age-related decline arises from the gradual accumulation of intracellular damages, e.g., genomic instability (3), protein misfolding and organelle dysfunction (4), that eventually overwhelm repair and protective systems (5). This damage-centered paradigm has profoundly shaped both the theoretical models of aging and the practical intervention strategies. Accordingly, major classes of anti-aging approaches, including metabolic interventions (e.g., dietary restriction, exercise, hormesis) (6–8), pharmacological targeting of conserved longevity pathways (e.g., mTOR inhibition, senolytics, NAD⁺ boosters) (9–11) and cellular reprogramming (12), aim to attenuate damage, enhance repair or activate cytoprotective responses. Despite their diversity, these strategies share a common conceptual orientation: they are fundamentally reactive, acting after deterioration has occurred (13). Implicit in this framework is a theoretical assumption that aging is primarily a problem of molecular-level deterioration rather than system performance (14). From a system perspective, this assumption imposes a fundamental limitation. Even with modest molecular damage, a biological system whose internal regulatory processes progressively lose efficiency in timing, coordination or energy utilisation, will inevitably suffer functional instability (15). In this regime, molecular damage is not only a cause of decline but also an emergent consequence of systemic inefficiency, creating a self-reinforcing cycle that damage-centered interventions can only partially offset.

An alternative, yet underexplored, perspective is that aging arises from a progressive loss of integrated cellular operation efficiency. Rather than being driven solely by damage accumulation, functional decline may reflect a reduced capacity of cells to process information, generate energy, maintain biogenesis, and preserve homoeostasis efficiently. This reframing shifts the emphasis from static molecular states to dynamic system performance and suggests that effective interventions would need to act on the dynamics of cellular processes themselves. Physical interventions are uniquely positioned in this regard, as they can, in principle, couple directly to system dynamics without relying on biochemical pathway specificity. In this context, mid-infrared (MIR) light has emerged as a particularly intriguing modality. Frequency-specific MIR stimulation has been shown to exert nonthermal and energy-efficient effects on neuronal signaling through resonance with ion-channel dynamics (16,17). MIR photons associated with biochemical reactions have been reported to influence DNA replication and assembly in model systems (18–20). Although these observations were not framed in the context of aging, they provide converging evidence that biological function can be modulated through frequency-matched physical interactions, pointing to a plausible route for influencing cellular efficiency at a systemic level.

In this study, we ask whether aging can be systemically prevented by enhancing cellular operation efficiency through physical interventions. To address this question, we analyze the aging process of *Caenorhabditis elegans* (*C. elegans*) across organismal, cellular and molecular scales under frequency-specific, super-weak MIR light. By using MIR light tuned to a narrow frequency window and delivered at super-low, nonthermal power, we examine whether a physical perturbation can simultaneously influence two central pillars of cellular performance: gene transcription and mitochondrial energy metabolism. This multiscale framework allows us to assess how frequency-matched physical inputs reshape aging trajectories and cellular homoeostasis, and to evaluate an efficiency-first mode of aging modulation at the level of the whole organism.

## Results

### Frequency-specific, super-weak MIR light modulates organismal aging in *C. elegans*

To examine whether a physical, nonthermal perturbation can produce an effect on organismal aging, we employed *C. elegans* as a genetically homogeneous and temporally well-resolved model system, in which aging trajectories can be quantitatively analyzed across the entire worm lifespan (21). To isolate nonthermal effects, we designed a MIR illumination scheme that delivered frequency-specific light (34.1 ± 0.1 THz, i.e., 8.79 ± 0.03 μm in wavelength, labelled 34-THz light) at a super-low average power density of ∼1 μW mm^−2^ across the culture dish using a single-sided frosted glass diffuser (Fig. 1*A*, more in *SI Appendix,* Fig. S1 and *Section 1*). Under these conditions, thermal imaging measurements showed that the temperature across the sample plane remained within 24.6–25.2 °C, closely matching the ambient temperature maintained at 25 °C (Fig. 1*A*), indicating negligible heating during MIR exposure. At this power level, the photon flux corresponds to a super-weak regime, with substantially fewer than one photon simultaneously traversing a single cell on average (*SI Appendix, Section 2*).

**Fig. 1.**
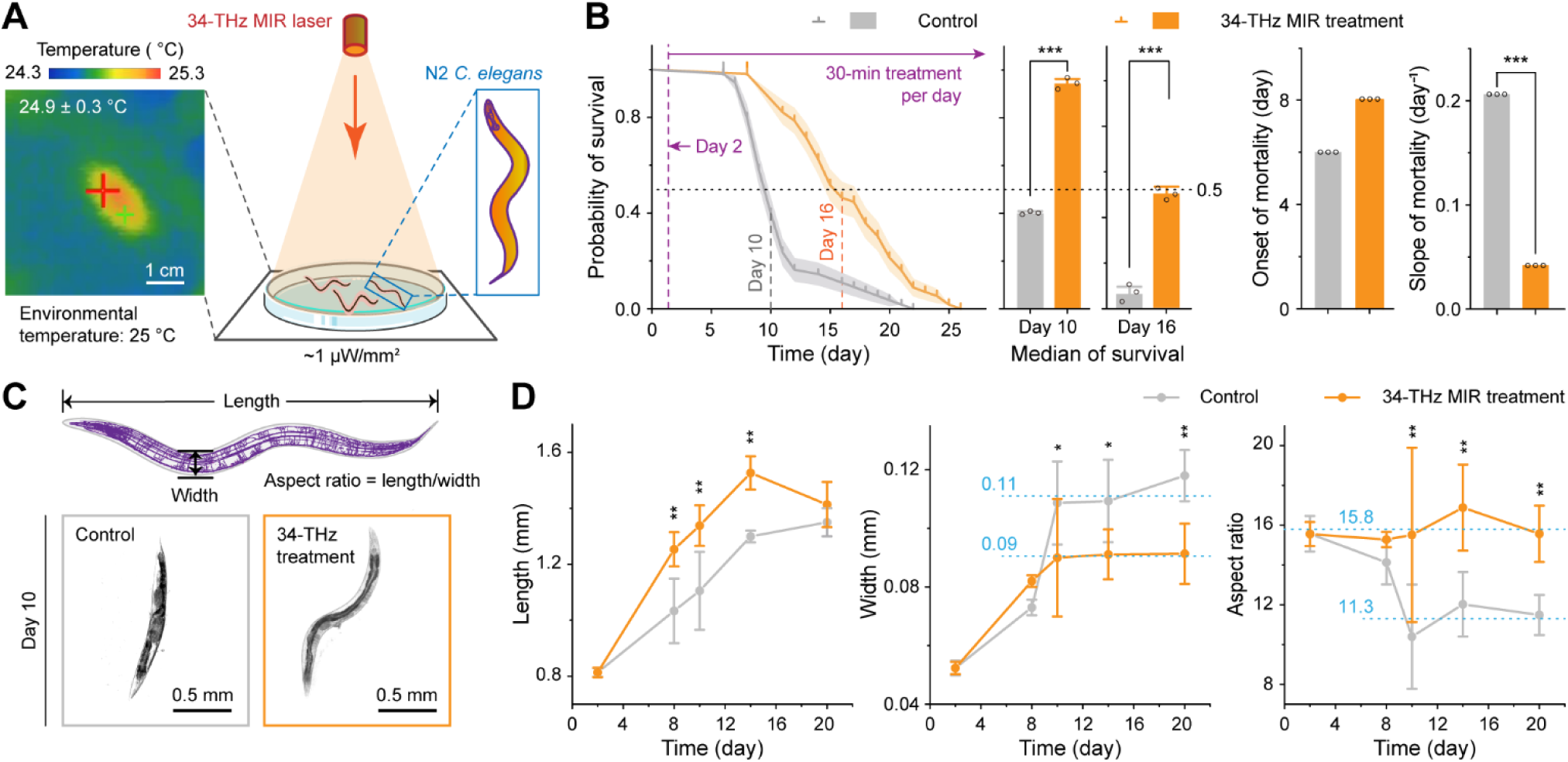
Effects of 34-THz MIR light on the *C. elegans* lifespan and morphology. **(*A*)** Schematic representation for MIR light modulating wild-type N2 *C. elegans*. Left: Worms in a culture dish treated by a laser with a mid-infrared (MIR) frequency of 34 THz and an intensity of ∼1 μW/mm^2^. Right: A two-dimensional thermal imaging of the culture-dish plane. The average temperature is 24.9 ± 0.3 °C. Scale bar: 1 cm. **(*B*)** Effects of 34-THz light on the worm lifespan. Left: Lifespan of the control (grey) and 34-THz MIR treated (orange) worms. The treatment begins at day 2 of adulthood. Middle: Survival probabilities at day 10 (control’s median of survival) and day 16 (treatment’s median of survival). Middle right: Onset and slope of mortality. **(*C*-*D*)** Effects of the 34-THz light on the *C. elegans* morphology during ageing. **(*C*)** Schematic representation of a worm (upper) and the representative bright-field micrographs of control (lower left) and 34-THz MIR treated (lower right) worms. Scale bar: 0.5 mm. **(*D*)** Effects of the 34-THz light on the worm length (left), width (middle) and aspect ratio (right) during ageing. Data are mean ± s.d., *n* = 3 (***B***) and 30 (***D***) groups, with 50–60 (***B***) and 1 (***D***) worms per group. ns, no significant difference (*P* > 0.05); *, significant difference (*P* < 0.05); ***, highly significant difference (*P* < 0.001).

Using this nonthermal MIR protocol, we applied the 34-THz light to *C. elegans* for 30 min per day beginning on day 2 of adulthood, a stage that precedes the onset of age-associated mortality in worms. Control populations were maintained under identical environmental conditions without MIR exposure. This experimental design enabled us to assess whether frequency-specific MIR light alters aging dynamics at the organismal level, independent of thermal or stress-related confounders. Before the experiments above, we first applied a commonly-used 808-nm infrared laser of ∼1 μW mm^−2^ to examine infrared thermal effects on the dynamics of our biological system, though those on the environmental temperature were already assessed. No significant changes during aging of worms caused by the light were observed (*P* > 0.05) (*n* = 3 groups, and 50–60 worms per group), indicating that the infrared thermal effects of this power on our aging dynamics can be ignored (*SI Appendix,* Fig. S2).

Survival analysis revealed a pronounced reshaping of aging trajectories under the 34-THz MIR light (Fig. 1*B*). At day 10 of adulthood, survival probabilities were 40 ± 7% for control worms and 97 ± 2% for MIR-treated worms (*P* < 0.001), and by day 16 these values declined to 11 ± 4% and 47 ± 7%, respectively (*P* < 0.001) (*n* = 3 groups, and 50–60 worms per group). The median lifespan increased from 10 days in the control group to 16 days in the treated group, corresponding to a 60% extension. This difference was statistically supported by a Gehan–Breslow–Wilcoxon test (*P* < 0.001) and a reduced hazard ratio (Mantel–Haenszel HR, 95% CI: 0.09–0.26).

Beyond extending lifespan, the MIR light markedly altered the temporal structure of aging. Analysis of survival dynamics showed that the onset of age-associated mortality occurred at day 6 in control worms but was delayed to day 8 in treated worms, representing a 33.3% delay in aging onset (Fig. 1*B*). More importantly, the decline in survival probability following onset was substantially attenuated under MIR treatment: the mortality slope decreased from 20.75 ± 0.05% per day in controls to 4.34 ± 0.05% per day in treated animals (*P* < 0.001) (*n* = 3 groups, and 50–60 worms per group), corresponding to a 4.8-fold deceleration of late-life mortality. These results indicate that super-weak 34-THz MIR light not only extends the lifespan but also suppresses the abrupt late-life mortality collapse characteristic of “cliff-edge” aging.

To assess whether these changes in survival dynamics were accompanied by preserved organismal integrity, we examined age-associated morphological metrics, including body length, body width and their aspect ratio (Fig. 1*C*). As shown in Fig. 1*D* (more in *SI Appendix,* Fig. S3), while body length increased in both groups during early adulthood, MIR-treated worms exhibited a faster increase before day 14, followed by convergence with controls at later ages (*n* = 30 worms). In contrast, body width diverged after day 8, increasing more rapidly in control worms but remaining restricted in treated worms. Consequently, the length-to-width aspect ratio, as a macroscopic indicator of morphological health (22), was maintained at a stable plateau of 15.8 in the treated group from day 2 to day 20, whereas it declined rapidly from 15.8 to 11.3 in controls by day 10 and remained low thereafter. Together, these morphological analyses indicate that 34-THz MIR light preserves youthful body geometry throughout aging, even at advanced ages.

### Regulation of mitochondrial structure and function by 34-THz MIR light during aging

To determine whether the reshaping of organismal aging trajectories is accompanied by preserved cellular energy systems, we examined mitochondrial integrity and function during worm aging. Mitochondria are central to cellular energy metabolism and homoeostasis, and age-associated mitochondrial deterioration is closely linked to declining cellular performance (23). We thus asked whether super-weak 34-THz MIR light influences the structural and functional state of mitochondria during aging in *C. elegans*.

To assess mitochondrial structural integrity, we analyzed mitochondrial morphology in the body wall muscle cells of SJ4103 (*zcIs14*(*myo-3::GFP*(*mit*))), in which mitochondria are fluorescently labelled with green fluorescent protein (GFP) (24). Mitochondrial fragmentation, which increases with age and is commonly used as a marker of mitochondrial dysfunction and cellular senescence (25), was quantified across the aging process. Representative confocal micrographs at day 14 revealed that mitochondria in control worms were substantially fragmented, whereas those in MIR-treated worms largely retained a tubular morphology (Fig. 2*A*, more in *SI Appendix,* Fig. S4), consistent with the distinct aging trajectories observed at the organismal level (Fig. 1*B*).

**Fig. 2.**
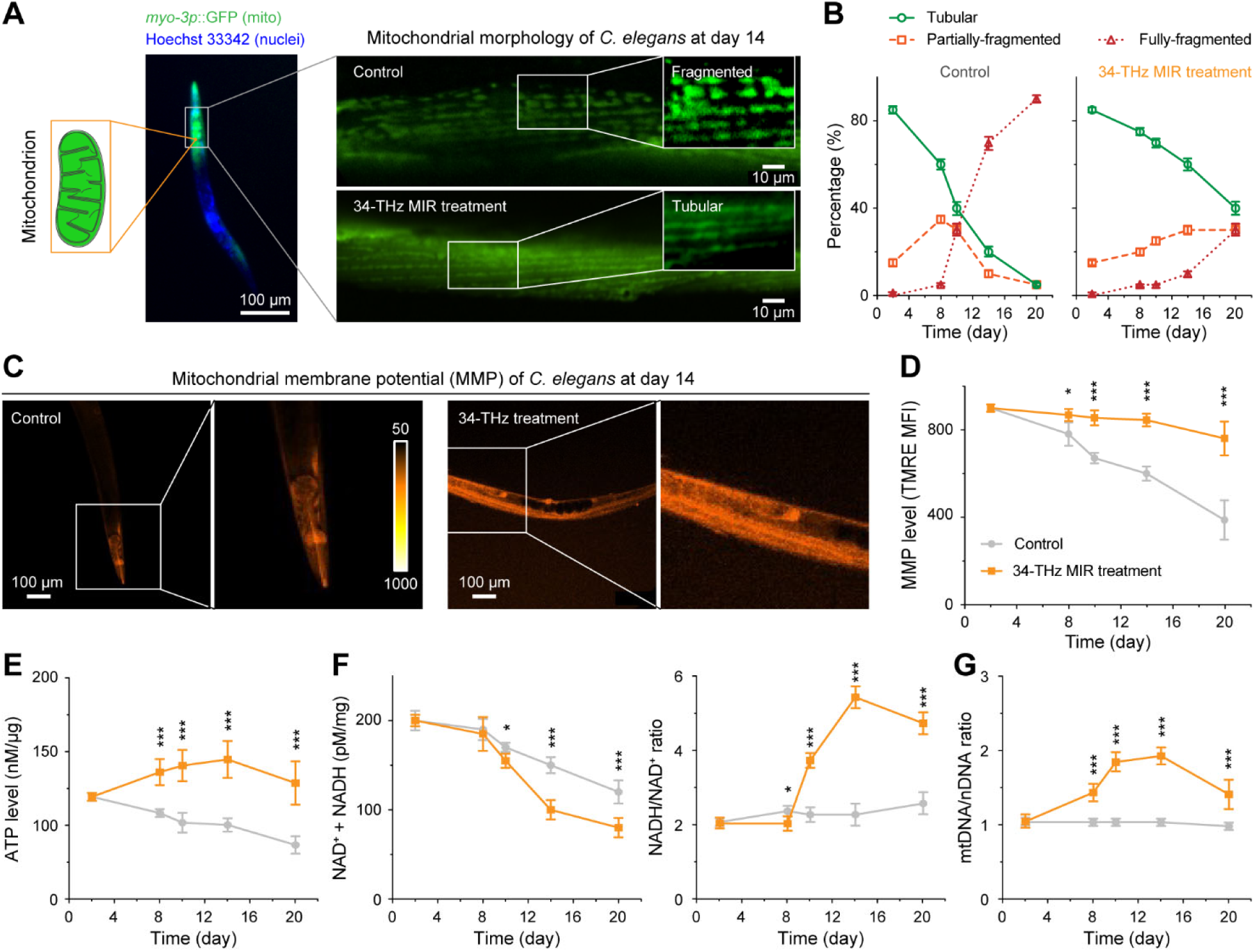
Effects of 34-THz MIR light on the mitochondria of *C. elegans* in the ageing process. **(*A*-*B*)** Mitochondrial morphology in the worm ageing process. **(*A*)** Representative fluorescence images of mitochondria in the worm at day 14. Fragmented mitochondria are clearly observed in the control group (upper), but not in the MIR treated group (lower). Scale bar: 100 µm for the main view and 10 µm for the zoom-in. **(*B*)** Morphological characteristics of mitochondria during ageing. According to morphology, mitochondria are divided to the tubular (solid green), partially-fragmented (dashed orange) and fully-fragmented (dotted red) ones. **(*C*-*D*)** Mitochondrial membrane potential (MMP) in worms during ageing. **(*C*)** Representative confocal fluorescence images of worm MMP at day 14. Color bar: fluorescence intensity. Scale bar: 100 µm for the main view and 50 µm for the zoom-in. **(*D*)** MMP levels of the control (grey) and 34-THz MIR treated (orange) groups in the ageing process. **(*E*-*G*)** ATP levels (***E***), NAD^+^ + NADH levels (***F***, left), NADH/NAD^+^ ratios (***F***, right) and mitochondrial DNA (mtDNA) copy numbers (***G***) of worms during ageing. Data are mean ± s.d., *n* = 3 (***B***), 30 (***D***) and 3 (***E*-*G***) groups, with 10 (***B***), 1 (***D***) and 300–350 (***E*-*G***) worms per group. ns, no significant difference (*P* > 0.05); *, significant difference (*P* < 0.05); ***, highly significant difference (*P* < 0.001).

Quantitative analysis showed that in control worms, the proportion of tubular mitochondria declined sharply from 85 ± 3% at day 2 of adulthood to 5 ± 3% at day 20 (*n* = 3 groups, 10 worms per group), accompanied by a corresponding increase in fully fragmented mitochondria from 0 ± 1% to 90 ± 3% (Fig. 2*B*). In contrast, MIR-treated worms maintained a substantially higher proportion of tubular mitochondria throughout aging, with 40 ± 6% tubular and only 30 ± 6% fully fragmented mitochondria remaining at day 20. These results indicate that the MIR treatment markedly preserves mitochondrial structural integrity during aging, even at advanced stages.

Following the preservation of mitochondrial structure, we next examined whether the MIR light maintains mitochondrial energetic function during aging. Mitochondrial membrane potential (MMP), which directly reflects the capacity for ATP synthesis, was measured in body wall muscle cells using the tetramethylrhodamine ethyl ester perchlorate (TMRE) probe (26). Representative fluorescence images at day 14 showed markedly stronger TMRE signals in MIR-treated worms than in controls (Fig. 2*C*, more in *SI Appendix,* Fig. S5), indicating an elevated MMP under MIR light. Quantitative analysis revealed a pronounced age-dependent decline in MMP of control worms, decreasing from 900 ± 7 at day 2 to 350 ± 37 at day 20 (*n* = 30 worms) (Fig. 2*D*). In contrast, MIR-treated worms exhibited only a modest reduction in MMP over the same period, from 900 ± 7 to 808 ± 25, indicating that 34-THz MIR light substantially prevents the age-associated loss of mitochondrial membrane potential, even at advanced ages.

To determine whether the preserved MMP translates into sustained energy output, we tested ATP levels during aging. At day 2, ATP concentrations were comparable between control and treated worms (∼120 nM μg^−1^) (*n* = 3 groups, 300–350 worms per group) (Fig. 2*E*). As aging progressed, ATP levels in control worms declined linearly to 86 ± 2 nM μg^−1^ by day 20. In contrast, MIR-treated worms exhibited an increase in ATP levels to 151 ± 2 nM μg^−1^ at day 14, followed by a moderate decrease to 135 ± 2 nM μg^−1^ at day 20, which remained significantly higher than the initial level at day 2 (120 ± 1 nM μg^−1^; *P* < 0.001). Together, these results indicate that 34-THz MIR light preserves mitochondrial energetic function during aging by sustaining both MMP level and ATP production.

To further assess whether the MIR light alters mitochondrial metabolic dynamics, we examined age-dependent changes in NAD(H)-related redox. The combined NADH + NAD⁺ level was used to estimate the total NAD(H) pool size, reflecting overall redox capacity, while the NADH/NAD⁺ ratio was used as an indicator of metabolic flux and redox state (27). As shown in Fig. 2f, for control worms, the NAD(H) pool size declined from 200 ± 5 pM mg^−1^ at day 2 to 120 ± 2 pM mg^−1^ at day 20 (*n* = 3 groups, 300–350 worms per group). In MIR-treated worms, this decline was more pronounced, reaching 80 ± 2 pM mg^−1^ at day 20, consistent with accelerated NAD(H) turnover under the MIR light. Despite this reduction in pool size, the NADH/NAD⁺ ratio in treated worms increased markedly during aging, rising from ∼2.0 in early adulthood to a peak of 5.5 ± 0.4 at day 14 and remaining elevated at 4.8 ± 0.4 at day 20. In contrast, the ratio in control worms remained relatively stable throughout aging. These results indicate that 34-THz MIR light enhances mitochondrial metabolic flux during aging, despite a reduced steady-state NAD(H) pool, consistent with a high-throughput, high-efficiency metabolic state.

Finally, to examine whether the enhanced mitochondrial function is accompanied by changes in mitochondrial abundance, we analyzed mitochondrial DNA (mtDNA) copy number along the aging trajectory. The copy numbers of mtDNA and nuclear DNA (nDNA) were quantified by quantitative polymerase chain reaction (qPCR), and the mtDNA/nDNA ratio was applied as an indicator of mitochondrial biogenesis (28). In control worms, the mtDNA/nDNA ratio remained approximately constant (∼1.0) throughout aging (*n* = 3 groups, 300–350 worms per group), meaning minimal changes in mitochondrial copy number (Fig. 2*G*). In contrast, MIR-treated worms exhibited a marked increase in mtDNA copy number, with the mtDNA/nDNA ratio rising from 1.04 ± 0.03 at day 2 to 1.85 ± 0.13 at day 10 and 1.93 ± 0.12 at day 14, before declining modestly to 1.41 ± 0.24 at day 20, remaining significantly higher than in controls. These results indicate that 34-THz MIR light promotes mitochondrial biogenesis during aging, contributing to the maintenance of mitochondrial system capacity even at advanced ages.

### Transcriptional regulation by 34-THz MIR light during aging

To determine whether the preserved mitochondrial system performance is accompanied by the coordinated regulation at the genetic level, we next examined transcriptional activity during aging. Gene transcription governs the flow of genetic information from DNA to RNA and plays a central role in maintaining cellular function and adaptability over time (29). We thus asked whether super-weak 34-THz MIR light modulates transcriptional programs associated with mitochondrial function and cellular defense during aging in *C. elegans*.

We first analyzed the transcription of mitochondrion-related genes during aging. In wild-type *C. elegans*, transcripts associated with mitochondrial biogenesis, energy production and protective responses exhibit only modest changes across the lifespan (*SI Appendix,* Fig. S6*A–C*) (30). In contrast, MIR-treated worms showed pronounced, age-dependent transcriptional modulation. The treated level of *hmg-5* relative to the untreated, as reflects mitochondrial DNA replication stability, increased during aging and reached a peak of 1.8 ± 0.2 at day 14 (*n* = 3 groups, 300–350 worms per group) (Fig. 3*A*). Similarly, the relative level of *nduo-1*, an important gene reflecting mitochondrial ATP production, increased to 1.5 ± 0.3 at day 14 before declining to 0.9 ± 0.2 at day 20, remaining close to baseline levels. These transcriptional patterns suggest that the MIR light counteracts the age-associated decline in mitochondrial biogenesis and bioenergetic gene transcription. In addition, the mitochondrial antioxidant gene *trx-2* exhibited a rapid increase to 2.9 ± 0.3 at day 14 (*n* = 3 groups, 300–350 worms per group), followed by sustained elevation at later ages (Fig. 3b), while the mitochondrial chaperone gene *hsp-6* also showed a pronounced mid-life peak of 2.6 ± 0.5 at day 14 (*n* = 3 groups, 300–350 worms per group) (Fig. 3*C*). Together, these results indicate that the 34-THz treatment engages transcriptional programs supporting mitochondrial maintenance and proteostasis during aging.

**Fig. 3.**
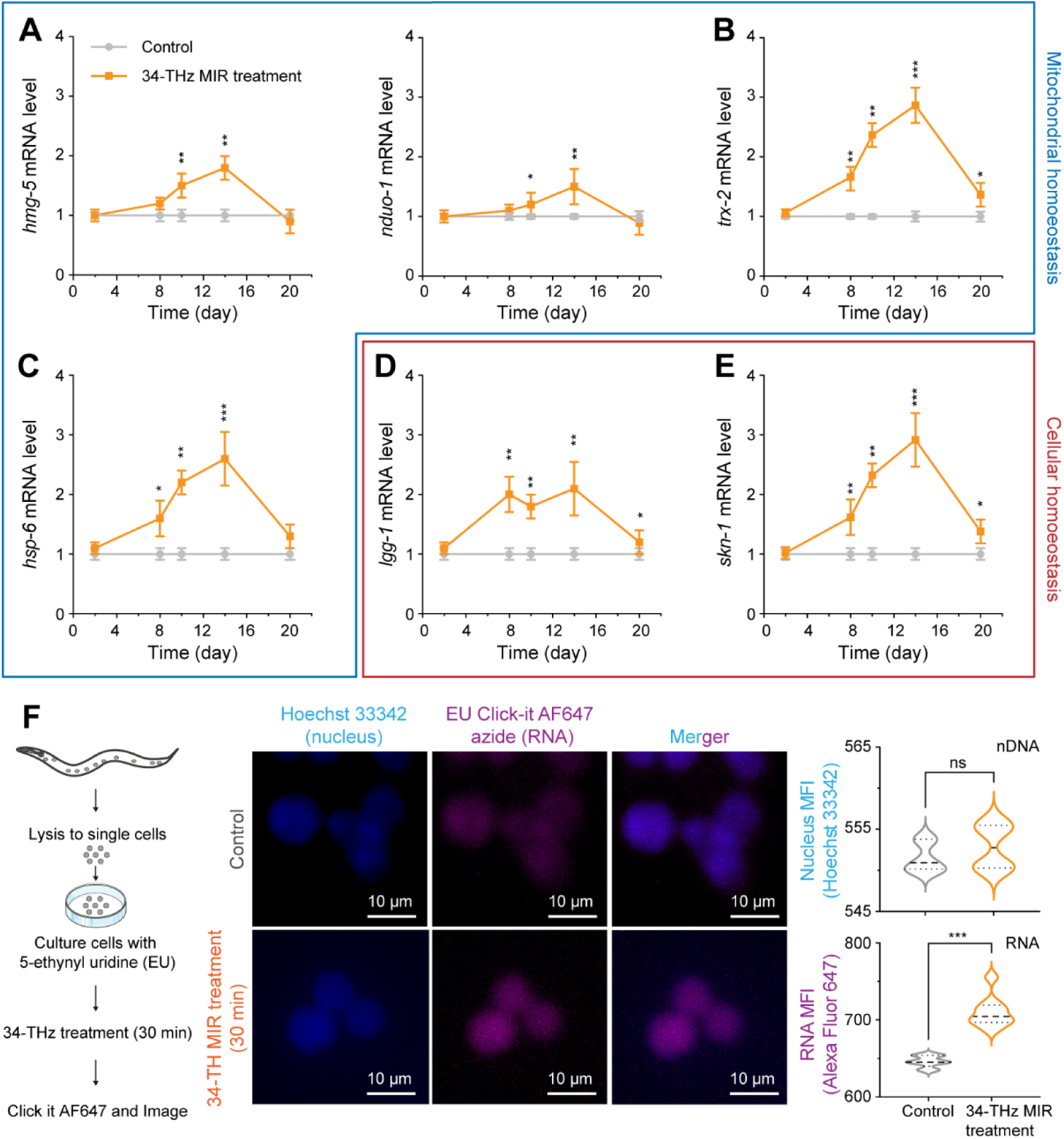
Effects of 34-THz MIR light on the ageing-related gene transcription in nuclei of *C. elegans* during ageing. **(*A*-*C*)** Gene transcription related to the mitochondrial homoeostasis regulation during ageing. **(*A*)** Mito biogenesis-related transcription. Left: Relative mRNA expression of *hmg-5*, related to the stability of mitochondrial gene replication. Right: Relative mRNA expression of *nduo-1*, related to mitochondrial complex I subunit involved in mitochondrial ATP production. **(*B*)** Mito redox homoeostasis-related mRNA expression of *trx-2*. **(*C*)** Mitochondrial unfolded protein response (UPR^mt^)-related mRNA expression of *hsp*-6. **(*D*-*E*)** Gene transcription related to the cellular homoeostasis during ageing. **(*D*)** Relative *lgg-1* mRNA level, related to the autophagy of cells. **(*E*)** Relative *skn-1* mRNA level, related to the redox homoeostasis of cells. **(*F*)** Effects of 34-THz light on the gene transcriptional activity. Left: A schematic diagram workflow for detecting newly synthesized RNA. Middle: Typical confocal fluorescence images of transcriptional activity in control and 34-THz MIR treated worm cells. Scale bar: 10 μm. Right: Median fluorescence intensity (MFI) of the nuclear DNA (Hoechst 33342 stained) and newly synthesized RNA (Alexa Fluor 647 azide stained) in the control (grey) and 34-THz MIR treated (orange) groups. In the panels ***A***-***E***: Data are mean ± s.d., *n* = 3 groups, 300–350 worms per group. ns, no significant difference (p > 0.05); *, significant difference (*P* < 0.05); **, very significant difference (*P* < 0.01); ***, highly significant difference (*P* < 0.001).

We next studied the effect of 34-THz MIR light on the transcription of genes related to the cellular defense. Given that LGG-1 is involved in the autophagy of cells (31) and SKN-1 is a key regulator of antioxidant defense that scavenges superoxide radicals (32), we employed the transcriptional level of their mRNA, *lgg-1* and *skn-1*, as aging biomarkers. In wild-type *C. elegans*, autophagic gene transcription reaches a peak in the mid-life, while the antioxidant gene response maintains a slight upregulation, followed by a decline in the late-life (*SI Appendix,* Fig. S6*D*,*E*) (30). The 34-THz MIR-treated levels of mRNA *lgg-1* and *skn-1* relative to the untreated are shown in Fig. 3*D*,*E*. We found that the relative level of *lgg-1* in MIR-treated worms rapidly increased to a ∼2.0 plateau from day 8 to 14, and then decreased to 1.2 ± 0.2 at day 20 (*n* = 3 groups, 300–350 worms per group). The relative level of *skn-1* rapidly increased from 1.0 ± 0.1 (day 2) to 2.9 ± 0.5 (day 14), and then decreased to 1.4 ± 0.2 (day 20) with a significant elevation compared to the initial level. These results indicate that the 34-THz MIR treatment upregulates the transcription of genes related to autophagy and antioxidant defense of cells during aging.

To investigate whether the 34-THz light modulates global RNA synthesis, we further quantified *de novo* RNA in light-treated and -untreated *C. elegans*. Single-cell suspensions were labelled with 5-ethynyl uridine (EU) analogue and treated with 34-THz MIR light for 30 min. Newly synthesized RNA was detected via click chemistry using Alexa Fluor 647 azide, with nDNA counterstained with Hoechst (33). The representative confocal fluorescence micrographs and the quantitative analyses of median fluorescence intensity (MFI) are shown in Fig. 3*F*. There was no significant difference in Hoechst fluorescence signal between the MIR-treated and -untreated groups, which reflects a stable experimental background and a consistent nDNA content, ensuring comparability and reliability of the subsequent RNA synthesis analysis. The Alexa Fluor 647 azide fluorescence intensity was markedly increased in the cells exposed to MIR light compared to the controls, denoting that the light enhances the synthesis rate of new RNA in the cells during the 30-minute period. Together, the 34-THz MIR light enhances the efficiency of *de novo RNA* synthesis in *C. elegans* cells, supporting its promotive effect on transcriptional activity.

### Regulation of cellular stress and responses by 34-THz MIR light during aging

To assess whether the enhanced efficiency of mitochondrial metabolism and gene transcription reduces the need for cellular stress compensation during aging, we next examined key markers of oxidative stress, cellular defense and longevity regulation. Oxidative stress, driven by the reactive oxygen species (ROS) accumulation, is a major contributor to age-associated cellular damage (34), and activation of antioxidant, autophagic and longevity pathways typically reflects an increased cellular stress burden (35). We therefore asked whether super-weak 34-THz MIR light alters the engagement of these stress-response systems during aging in *C. elegans*.

We first quantified intracellular ROS levels using the DCFDA probe and monitored antioxidant defense through SKN-1::GFP reporter worms (36,37). As shown in Fig. 4*A* (more in *SI Appendix,* Figs. S7,S8), at day 14, strong DCFDA fluorescence was observed in control worms, whereas MIR-treated worms exhibited minimal ROS signals. Across aging from day 2 to 20, ROS levels in control worms increased markedly from 100 ± 8 to 680 ± 72 (*n* = 3 groups, 10 worms per group), whereas MIR-treated worms showed only a modest increase from 97 ± 5 to 181 ± 53. In parallel, SKN-1 protein levels, which rose substantially with age in control worms, remained close to basal levels in treated ones throughout aging. These results indicate that the 34-THz MIR treatment suppresses age-associated oxidative stress accumulation, thereby reducing the requirement for antioxidant defense activation.

**Fig. 4.**
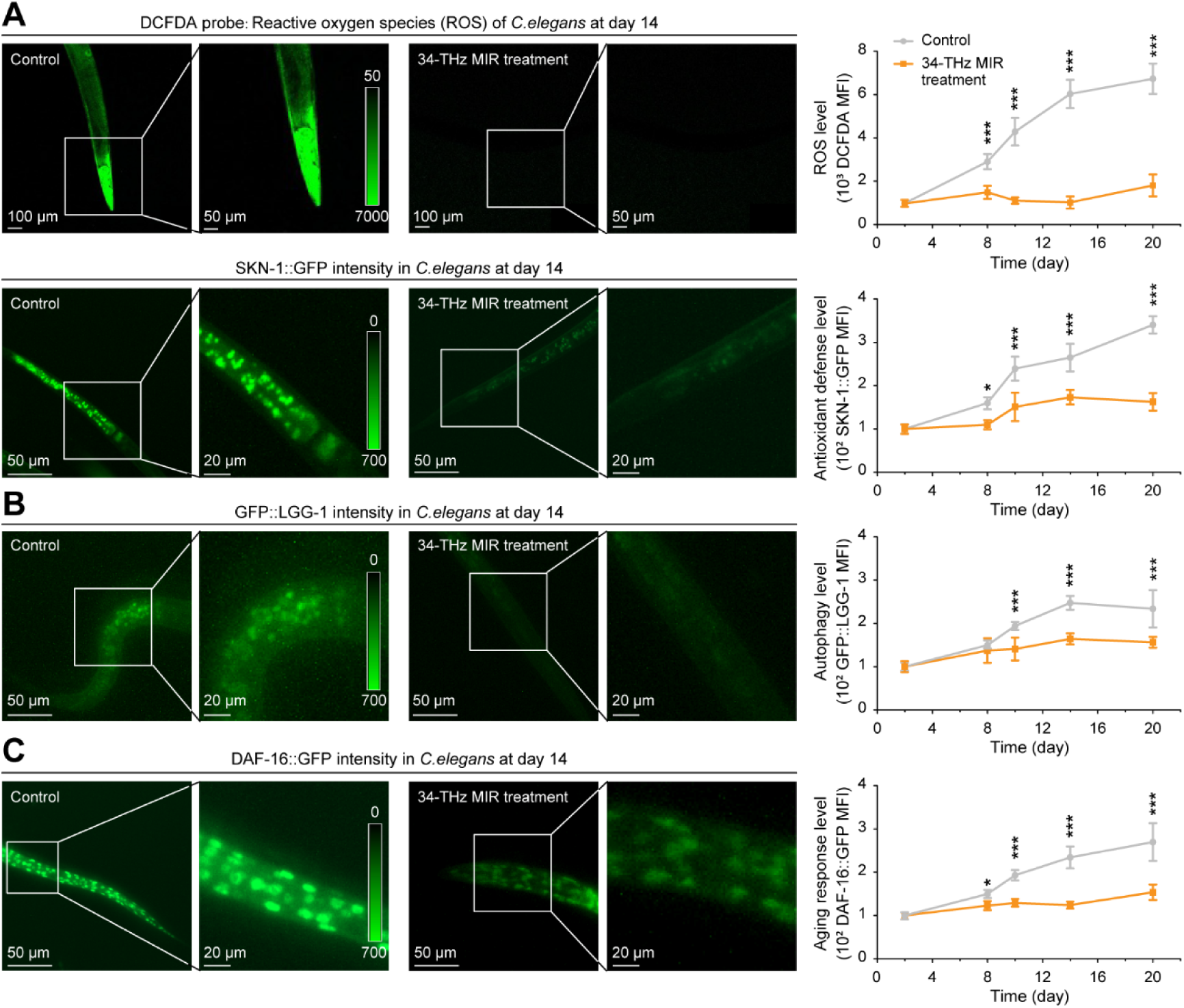
Effects of 34-THz MIR light on the cellular homoeostasis of *C. elegans* during ageing. **(*A*)** Effects of 34-THz MIR light on the oxidative stress regulation. Upper: Reactive oxygen species (ROS) levels of worms. Upper left: Representative confocal images. Color bar: fluorescence intensity. Scale bar: 100 μm for the main view and 50 μm for the zoom-in. Upper right: ROS levels of worms during ageing. The grey and orange curves indicate the control and 34-THz MIR treated groups, respectively. Lower: Antioxidant defense levels of SKN-1::GFP strain during ageing. Lower left: Representative fluorescence images. Scale bars: 50 μm for the main view and 20 μm for the zoom-in. Lower right: DAF-16::GFP levels during ageing. Lower right: SKN-1::GFP fluorescence levels during ageing. **(*B*)** Autophagy levels of GFP::LGG-1 strain. Left: Representative fluorescence images. Right: GFP::LGG-1 levels in worms during ageing. **(*C*)** Ageing response levels of DAF-16::GFP strain. Left: Representative fluorescence images. Right: DAF-16::GFP levels in worms during ageing. Data are mean ± s.d., *n* = 3 groups, 10 worms per group. ns, no significant difference (*P* > 0.05); *, significant difference (*P* < 0.05); ***, highly significant difference (*P* < 0.001).

We next studied the effect of 34-THz MIR light on the autophagic activity and longevity regulator in the cells of aged *C. elegans*. The autophagic activity and longevity regulator are usually marked by LGG-1 and DAF-16, respectively (38). LGG-1 and DAF-16 levels were measured by GFP::LGG-1 (39) and DAF-16::GFP (40) fluorescence worms, respectively. As the representative confocal images shown in Fig. 4b (more in *SI Appendix,* Fig. S9), compared to the control at day 14, the LGG-1 response was substantially suppressed in the MIR-treated group. During aging from day 2 to 20, the MFIs of control and treated groups significantly increased from 100 ± 3 to 234 ± 43 and slightly increased from 100 ± 3 to 157 ± 13 (*n* = 3 groups, 10 worms per group), respectively (Fig. 4*B*, right). As for the longevity regulator, at day 14, fluorescence in the control was much stronger than that in the treated group (Fig. 4*C*, more in *SI Appendix,* Fig. S10). From day 2 to 20, the control and treated MFIs significantly increased from 100 ± 3 to 270 ± 144 and slightly increased from 100 ± 3 to 154 ± 18 (*n* = 3 groups, 10 worms per group), respectively (Fig. 4*C*, right). These results indicate that the 34-THz MIR treatment maintains the cellular environment healthy and non-toxic close to the youthful state in the aging process of *C. elegans*.

In all, 34-THz MIR light effectively alleviates cellular stress by suppressing ROS accumulation, as in turn reduces the need for the organism to activate its key antioxidant defense (e.g., SKN-1), repair pathway (e.g., LGG-1) and even longevity regulation (e.g., DAF-16), which together preserve cellular homoeostasis close to the youthful basal level during aging.

## Discussion

The above results show that super-weak 34-THz MIR light can exert robust effects on aging in *C. elegans* across multiple biological scales. Together, these observations establish that a physical, nonthermal perturbation is capable of influencing long-term aging dynamics in a living organism (Fig. 5*A*). However, these findings also raise an important question: how can such a weak physical input produce sustained benefits without engaging classical damage-repair or stress-response pathways? Addressing this question requires moving beyond a damage-centered interpretation of aging and instead considering aging as a problem of declining system efficiency and performance. In the following discussion, we interpret our results within an efficiency-first framework and consider how the MIR light may act on core cellular processes to stabilize aging systems over time.

**Fig. 5.**
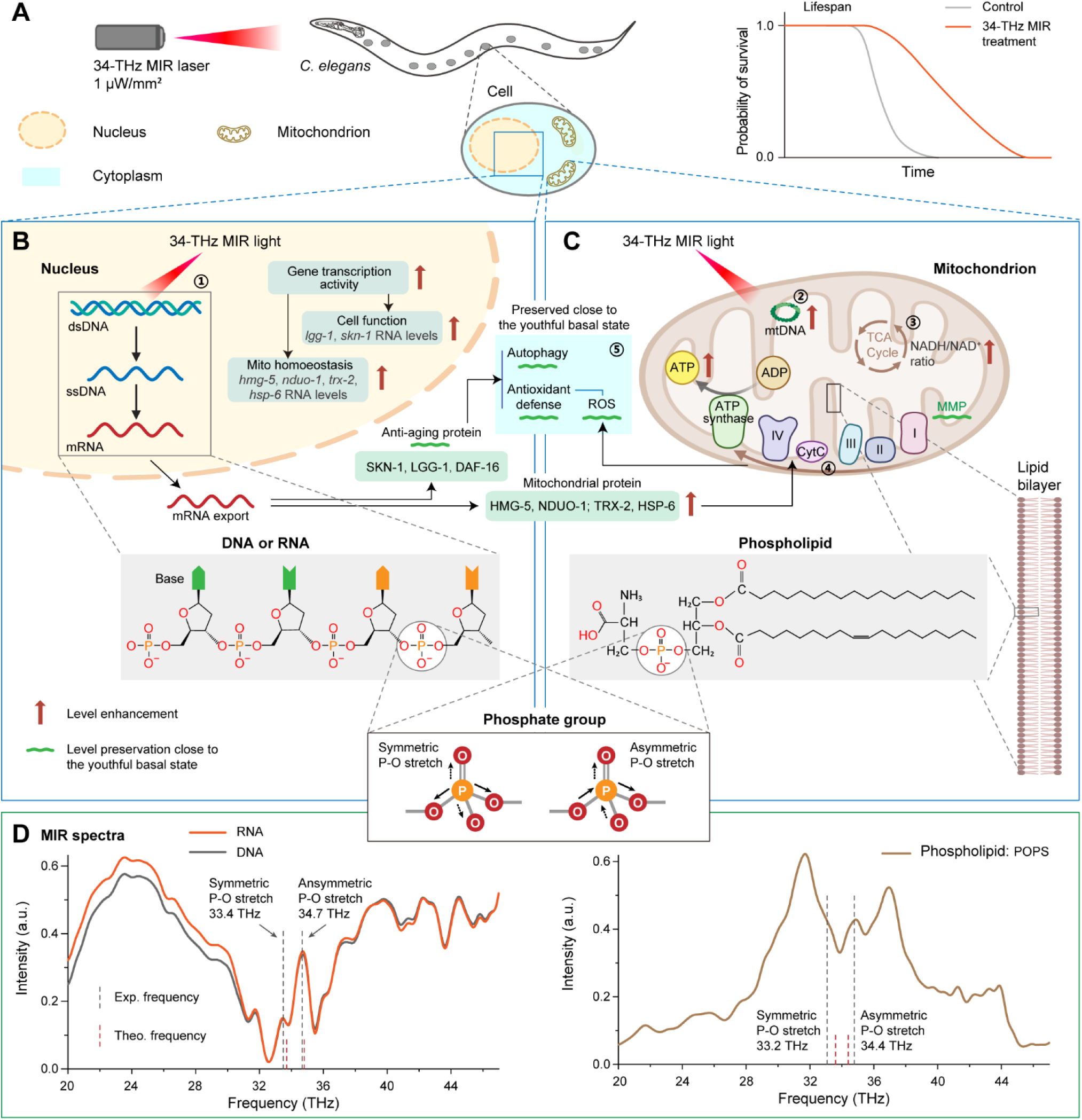
Summary of 34-THz light anti-ageing effects and the MIR spectroscopy analysis for the underlying mechanism. **(*A*-*C*)** Summary of the anti-ageing effects of 34-THz MIR light on *C. elegans*. **(*A*)** Effect on the lifespan of worms. **(*B*)** Effects on the gene transcription and protein expression. Compared to the untreated worms, the MIR light increases the overall efficiency of RNA transcription, which upregulates transcriptional levels of genes associated with cellular function (*lgg-1*, *sod-3*) and mitochondrial homoeostasis (*hmg-5*, *nudo-1*, *trx-2*, *hsp6*), maintains key factors of stress responses (SKN-1, DAF-16) and autophagy (LGG-1) on the levels close to those in the youthful state. **(*C*)** Effects on the mitochondria. The light enhances integrated function of mitochondria, i.e., clearly inhibiting the mitochondrial membrane potential (MMP) decrease, greatly elevating ATP production. The MIR light significantly upregulates the NADH/NAD^+^ ratio, suggesting a more efficient metabolic state. The copy number of mitochondrial DNA (mtDNA) is enhanced too. Note: The light modulation not only maintains the mitochondrial operation with a low ROS level, but also equips the cytoplasm with antioxidant proteins to neutralize ROS, preserving the cells to the youthful basal. The labels ① to ⑤ mean the cross-talk between nucleus (①) and mitochondria (②, ③, ④), and its results in cytoplasm (⑤) under the 34-THz MIR light. **(*D*)** MIR spectra of nucleic acid (left) and phospholipid (right) molecules. Two characteristic peaks in the range of 33–35 THz occur, which correspond to the symmetric and asymmetric stretches of P-O bonds in the phosphate group of nucleic acids and phospholipids.

Interpreted at the system level, our results indicate that the anti-aging effects of 34-THz MIR light arise from the coordinated maintenance of core cellular processes during aging, but not from the modulation of a single signaling pathway (Fig. 5*B*,*C*). In particular, MIR exposure is associated with a functional alignment between the transcriptional activity and the mitochondrial bioenergetic performance, while these two processes commonly become decoupled as organisms age (41). At the nuclear level (location ① in Fig. 5*B*), MIR treatment is accompanied by elevated transcriptional efficiency and increased RNA output, supporting the sustained expression of genes involved in mitochondrial biogenesis, cellular maintenance and stress response. Notably, these transcriptional changes occur with key regulatory factors remaining close to youthful basal levels of homoeostasis, but not sustained activation of classical stress-response pathways. In parallel, at the mitochondrial level (location ② in Fig. 5*C*), MIR exposure is associated with enhanced mitochondrial mtDNA abundance, accompanied by the preservation of mitochondrial structure and function, as reflected by maintained structural integrity, preserved MMP level and elevated ATP production. These observations define a coordinated cellular state in which transcriptional demand and mitochondrial energy supply remain mutually matched over time. This mode of regulation contrasts with lifespan-extending interventions that act by suppressing metabolism or inducing compensatory stress responses. Instead, frequency-specific MIR light appears to support the efficient operation of existing cellular machinery, thereby sustaining overall system performance during aging.

This coordinated state raises a key metabolic question: how can mitochondria sustain high energetic output while cellular stress remains low during aging? In MIR-treated animals, we observe an unusual combination, which are a smaller total NAD(H) pool alongside a higher NADH/NAD⁺ ratio in the tricarboxylic acid (TCA) cycle (Fig. 2*F*, summarized in the location ③ of Fig. 5*C*). In the conventional view, an elevated NADH/NAD⁺ ratio is usually associated with metabolic stress or impaired electron transport chain (ETC) function (42). However, this view is not supported by our functional measurements. Despite the elevated NADH/NAD⁺ ratio, MIR-treated worms maintain MMP levels and exhibit higher ATP levels (location ④ in Fig. 5*C*), indicating that electron transport remains efficiently coupled to ATP synthesis rather than compromised. These observations suggest that MIR exposure alters the mode of mitochondrial energy metabolism, not by expanding redox capacity, but by reshaping TCA and ETC flux dynamics. One plausible interpretation is a high-turnover, “just-in-time” metabolic regime. In this mode, reducing equivalents are generated and consumed more efficiently and rapidly, enabling high flux with a smaller steady-state NAD(H) pool while sustaining energetic output. Such kinetic efficiency provides a coherent explanation for the simultaneous preservation of bioenergetic performance and the reduced requirement for stress-compensatory responses observed under the 34-THz MIR light, reinforcing the view that aging modulation in this system arises from improved operational efficiency rather than damage mitigation.

If mitochondrial energy metabolism operates with higher kinetic efficiency, a direct consequence should be a reduced generation of cellular stress. Consistent with this expectation, MIR-treated worms show markedly lower accumulation of reactive oxygen species during aging, together with diminished engagement of antioxidant, autophagic and longevity-associated responses (Fig. 4, summarized in the location ⑤ of Fig. 5*B*,*C*). Importantly, this pattern does not indicate a suppression of cellular defense mechanisms. Instead, the preserved mitochondrial function and efficient electron transport limit oxidative burden at its source, thereby reducing the requirement for downstream stress compensation. As a result, regulators (e.g., SKN-1, LGG-1 and DAF-16) remain close to youthful basal levels, reflecting a stable, low-stress operating state rather than impaired protective capacity. These observations suggest that efficiency-first modulation stabilizes aging cells by preventing the emergence of stress signals that would otherwise trigger chronic defense activation. In this view, low stress is not an active target of intervention, but an emergent property of sustained system efficiency, i.e., a state with the reinforced integrity of mitochondrial-nuclear cross-talk that maintains youthful organelle coordination and pre-empts chronic compensatory responses.

Last, a key remaining question is how this super-weak frequency-specific MIR light input can concurrently influence transcriptional dynamics and mitochondrial bioenergetics without invoking biochemical pathway specificity. To address this question, we sought molecular-level evidence for a shared physical susceptibility in core cellular components. Using Fourier transform infrared (FT-IR) spectroscopy, we found that purified DNA and RNA exhibited two prominent absorption peaks in the 33–35 THz window (particularly 33.4 and 34.7 THz), and that a representative phospholipid POPS showed closely aligned peaks (33.2 and 34.8 THz) within the same frequency range (Fig. 5*D*). The density functional theory (DFT) calculations further assigned these features to the symmetric and antisymmetric P–O stretching modes of phosphate groups (PO_4_) in nucleic acids and phospholipids, consistent with the previous reports (43,44). These results provide a molecular context consistent with the dual-system effects summarized in Fig. 5*B* and *C*. Phosphate groups are ubiquitous at the interfaces that govern information processing and energy transduction. In nucleic acids, they shape the electrostatic and mechanical environment that constrains template accessibility and polymerase progression; in phospholipid bilayers, they contribute to the physical properties of mitochondrial membranes that support the cristae organization and the function of embedded enzymes. From this view, MIR light tuned to the PO_4_ vibrational window could modulate cellular performance by subtly shifting the dynamic properties such as conformations of these phosphate-rich macro-biomolecular frameworks, thereby influencing transcriptional kinetics and mitochondrial membrane-associated processes in parallel.

We emphasize that resonance alone does not establish a complete mechanistic chain from photon absorption to organismal lifespan extension, and additional work will be required to resolve how vibrational excitation propagates across molecular, organellar and cellular scales. Nevertheless, the convergence of (i) system-level preservation of transcriptional activity, mitochondrial function and cellular low-stress state, and (ii) frequency-aligned phosphate vibrational modes in nucleic acids and phospholipids, supports an efficiency-first model in which physical, frequency-matched inputs can enhance the kinetic performance of existing cellular systems. More broadly, this framework suggests a complementary route for aging modulation, as it tunes the operational efficiency of core processes rather than compensating for accumulated damage or inducing chronic stress responses.

In summary, by analyzing aging across organismal, cellular and molecular scales in *C. elegans*, we demonstrate that super-weak, frequency-specific MIR light can systemically modulate the aging process in a nonthermal manner. Exposure to 34-THz MIR light robustly prolongs lifespan, delays the onset of age-associated decline and preserves cellular homoeostasis into advanced age. These effects arise from the enhanced coordination and efficiency of core cellular processes, including gene transcription and mitochondrial metabolism, rather than the activation of damage-repair or stress-response pathways. At the molecular level, the alignment of MIR frequency with phosphate vibrational modes shared by nucleic acids and phospholipids provides a plausible physical basis for simultaneously regulating information processing and energy transduction. This insight suggests a general principle for aging modulation: physical interventions tuned to intrinsic frequency windows of molecular dynamics may enhance the dynamic performance of existing cellular systems, rather than introducing new biochemical components or pathways. More broadly, our findings highlight frequency-specific, nonthermal physical modulation as a framework for exploring efficiency-first strategies to stabilize biological function during aging, offering a complementary direction for future anti-aging research beyond conventional damage-centered approaches, such as advanced clinical phototherapies and integrated wearable wellness devices.

## Materials and Methods

### *C. elegans* strains and maintenance

*Caenorhabditis elegans* strains were obtained from the Caenorhabditis Genetics Center (CGC, University of Minnesota): N2 (Bristol wild-type) (**21**), SJ4103 [zcIs14(myo-3::GFP(mit))] (24), LG333 [skn-1::GFP] (37), DA2123 [adIs2122(lgg-1p::GFP::lgg-1)] (39), and TJ356 [zIs356(daf-16p::daf-16a/b::GFP)] (40). All strains were verified for genotype by PCR and phenotype by morphological inspection. Transgenic strains were backcrossed against N2 background (4–6 generations) to minimize genetic drift. Worms were cultured at 20 ± 0.2 °C on NGM plates with a two-layer agar system: 15 mL NGM (2% agar) bottom layer and 25 mL NGM (3% agarose) top layer, both supplemented with gentamicin (16.7 μg mL^−1^) and carbenicillin (25 μg mL^−1^). OP50 (cultured to OD_600_ ≈ 1.5) was killed with gentamicin (200 μg mL^−1^, 5 h), concentrated 75-fold, and seeded onto plates to provide consistent food while preventing bacterial proliferation.

### Worm synchronization

Synchronized L1 larvae were generated by bleaching gravid adults (18–24 h post-L4) with alkaline hypochlorite (0.5 M KOH, 10% NaClO) for 5 min with visual microscopic monitoring. Embryos were washed three times in M9 buffer and hatched overnight at 20 °C with agitation (180 rpm) to reach L1 arrest. L4-stage larvae (40–42 h development; confirmed by morphological inspection) were used as the starting population for all experiments.

### 34-THz MIR light exposure system

The laser system was configured vertically with a total distance of 32 cm from the output aperture of MIR source (continues-wave quantum cascade laser, 34.1 ± 0.1 THz, i.e., 8.79 ± 0.03 μm in wavelength) to the sample plane. A single-sided ground glass diffuser was integrated into the optical path, positioned 27 cm downstream from the laser exit and 5 cm above the sample plane. This configuration served two purposes: (1) expanding the beam to achieve uniform coverage across 30-mm culture dishes, and (2) providing effective thermal isolation between the laser source and the biological sample.

The incident MIR optical power at the sample plane was directly measured using a calibrated mid-infrared laser power meter (Ophir VEGA, Ophir Optronics Solutions Ltd., Israel; wavelength range 0.19– 20 µm) with the culture-dish lid removed. Irradiance was calculated from the measured power and the effective beam area determined from beam expansion geometry. Under experimental conditions, MIR light was maintained at approximately 1 μW mm^−2^. The ground glass diffuser homogenized the MIR light, minimizing hotspots and ensuring uniform illumination across the culture dish.

To verify the biological effects specifically induced by 34-THz light resonance rather than photothermal artifacts, comprehensive thermal analysis of the optical setup was performed using a thermos imaging camera (Hikvision DS-2TP31-3AUF Handheld Thermography Camera). Real-time thermographic imaging was conducted to monitor temperatures at the diffuser surface and at the culture-dish interface during irradiation. The spatial separation (5 cm air gap) between the diffuser and sample plane was designed to thermally isolate the culture area from the laser source, ensuring that worms experienced a nonthermal MIR electromagnetic field. No detectable temperature increase was observed at the sample plane during MIR exposure.

### Daily MIR light exposure

Continuous 30-min exposure per day. Control groups were maintained under identical environmental conditions without laser exposure.

### Lifespan assays

Lifespan analyses were initiated at day 2 of adulthood. Worms were exposed to daily 30-min MIR treatment (or identical conditions without MIR light for the control groups). Survival was monitored at 24-h intervals by gentle stimulation with a platinum wire and non-responsive worms were scored as dead and removed immediately. Minimum *n* = 50–60 worms per condition, three independent biological replicates per strain. Survival data were plotted as Kaplan–Meier curves and statistically evaluated using the log-rank (Mantel–Cox) test.

### Morphological analysis

Brightfield images were acquired at day 2, 8, 10, 14 and 20 of adulthood using a Thermo EVOS M300 imaging system (20× objective; 2,048 × 2,048 pixels, spatial resolution 0.5 μm per pixel). Quantitative image analysis was performed using ImageJ/Fiji (version 1.53, NIH). For each image, automated segmentation was conducted: worm bodies were identified using Otsu’s binary thresholding method, followed by morphological operations (dilation and erosion) to refine body boundaries. An ellipse was fitted to the segmented worm body using the Fit Ellipse function to extract body length (major axis), body width (minor axis), and aspect ratio (length/width). Measurements were normalized to each worm’s baseline of day 0 to account for initial size variation. At each timepoint, 20– 30 worms per condition were analyzed (22).

### Mitochondrial morphology analysis

At day 14 of adulthood, SJ4103 worms expressing mitochondrially targeted GFP in body wall muscle were immobilized in 10 mM levamisole and imaged using a Zeiss LSM900 confocal microscope (488-nm laser, 5–8% power, 1 AU pinhole) [24]. Mitochondrial networks in body wall muscle were classified as: tubular (>80% connectivity), partially fragmented (30–80%), or fully fragmented (<30%). Mean fluorescence intensity was quantified using ImageJ/Fiji (version 1.53) with background subtraction from regions devoid of muscle tissue. Morphological classification was performed by an investigator blinded to experimental conditions.

### Mitochondrial function assessment

The MMP was assessed using TMRE (100 nM, 1 h, 20 °C; Sigma-Aldrich T669) (26), and reactive oxygen species (ROS) levels using DCFDA (10 μM, 2 h incubation at 20 °C; Thermo Fisher Scientific, D399) (36). Both stained worms were imaged using a Zeiss LSM900 confocal microscope (488-nm laser for DCFDA, 561-nm laser for TMRE; 5–8% power, 1 AU pinhole) or Thermo EVOS M300 imaging system. Quantification of mean fluorescence intensity was performed using ImageJ/Fiji (version 1.53) after background subtraction.

### Biochemical measurements

Approximately 100 worms were homogenized in ice-cold lysis buffer (MCE HY-K0314-100T), centrifuged (12,000 × g, 10 min, 4 °C), and ATP quantified immediately by luciferase assay (Promega L0610) using SpectraMax iD5 microplate reader. ATP content was normalized to total protein concentration, determined using BCA assay (Pierce, 23225).

Approximately 2,000 worms were extracted (Abcam ab65348). Total NAD was measured at 450 nm using SpectraMax iD5; NADH was measured after heating at 60 °C for 30 min (selective NAD⁺ degradation). NAD⁺ = Total NAD – NADH (**27**).

Worms (∼200) were dissociated into single cells using chitinase (2 mg mL^−1^, Sigma C6137) and pronase (5 mg mL^−1^, Sigma 10165921001; 37 °C, 20 min). Cells were incubated in 1 mM 5-EU (Jena Bioscience EU-A815), exposed to 30-min THz MIR light, fixed (4% PFA), permeabilized (0.5% Triton X-100), and labeled via click chemistry (Thermo Fisher C10642). Hoechst 33342 (1 μg mL^−1^) stained nuclei [33]. Fluorescence microscope imaging quantified AF647-to-Hoechst fluorescence ratio per cell as a transcription index using ImageJ/Fiji (version 1.53) with background subtraction (*n* = 50–100 cells per sample).

### Gene expression analysis

Total RNA from approximately 100 worms was extracted using TRIzol (Tiangen 9108Q), reverse-transcribed (Bimake B24408M), and analysed by real-time quantitative polymerase chain reaction (*RT*-*qPCR*) using SYBR Green Master Mix on Roche LightCycler 480 system. All primers were validated: A range of 90–110% for the amplification efficiency and an R^2^ value of 0.98 were considered acceptable, single melt peaks, and correct amplicon size on agarose gels. Reactions (20 μL; 250 nM primers, 1:10 diluted cDNA, 10 μL SYBR Green) ran in technical triplicate with no-template controls. Data were analysed using the ΔΔCt method with *act-1* as the internal reference gene. Genes analysed: *nduo-1*, *hmg-5*, *hsp-6*, *trx-2*, *lgg-1*, *skn-1*, *cox-1*. Single-worm genomic DNA (Proteinase K lysis) was quantified by RT-qPCR as the *cox-1*/*act-1* ratio (both primers: 95–110% efficiency) (*SI Appendix,* Table S1) (28).

### Longitudinal fluorescence imaging of stress-response reporters

Transgenic *C. elegans* strains expressing stress-responsive fluorescent reporters (SKN-1::GFP, GFP::LGG-1, DAF-16::GFP) were exposed to daily 30-min THz MIR light and imaged on days 2, 8, 10, 14, and 20 of adulthood to assess age-dependent regulation of cellular stress responses. Fluorescence imaging on days 2, 8, 10, and 20 was performed using an Evos M3000 (Thermo Fisher) widefield fluorescence microscope with automated image acquisition. At each timepoint, 20–30 worms per condition were imaged. GFP fluorescence intensity were quantified in a standardized anterior body wall region using ImageJ/Fiji (version 1.53) with background subtraction.

### Fourier transform infrared spectroscopy (FT-IR) analysis

FT-IR in attenuated total reflection mode (ATR) was used to assess vibrational features of nucleic acids and phospholipids. Pure powder samples of nucleic acids (DNA, RNA; Sigma-Aldrich) and phospholipids (POPS, Avanti Polar Lipids) were used as model molecules. Approximately 2–3 mg of each sample was directly placed onto a diamond attenuated total reflection (ATR) crystal plate.

FT-IR ATR spectra were acquired using a Bruker TENSOR 27 Fourier Transform Infrared Spectrometer equipped with a liquid nitrogen-cooled mercury cadmium telluride (MCT) detector. Prior to spectral acquisition, the instrument was purged with dry nitrogen gas and cooled with liquid nitrogen to ensure optimal signal-to-noise ratio and spectral stability. Measurements were collected in the wavenumber range of 4000–600 cm^−1^ at 4 cm^−1^ resolution with 32 co-added scans per sample. Baseline correction and spectral processing were performed using Bruker OPUS software (version 8.0). For each powder sample type (DNA, RNA, POPS), three independent measurements were conducted.

### Statistical analysis

All analyses used GraphPad Prism 10.1.1. The data are presented as the mean ± s.d. (standard deviation). Data normality was assessed by D’Agostino-Pearson test; variance homogeneity by Levene’s test. Kaplan–Meier survival curves were compared using Log-rank (Mantel– Cox) test, reporting χ² and p-value. Multi-group comparisons employed one-way ANOVA (or Kruskal-Wallis for non-normal data) followed by Tukey’s post-hoc test (family-wise error: 0.05). Pairwise comparisons used two-tailed unpaired t-tests (Welch’s correction) or Mann-Whitney tests, with effect sizes (Cohen’s d, hazard ratios with 95% CI) reported. All figures were assembled using Adobe Illustrator 2020 following Nature guidelines (line weight 0.5–1.0 pt, Arial font 8–12 pt, color-blind-friendly palettes).

### Density functional theory calculations

Density functional theory (DFT) calculations were performed using the M06-2X exchange-correlation functional in conjunction with the 6-31G(d) basis set. Solvation effects were modeled using the polarized continuum model (PCM), with physiological solution treated as the solvent. Vibrational frequency calculations were carried out to obtain molecular vibrational modes. The frequencies of photon absorption and emission were identified from vibrational modes in which the PO_4_ functional group exhibits significant atomic displacements and high infrared intensities. All DFT calculations were performed using the Gaussian 16 software package.

## Author Contributions

Q.H. and B.S. conceived the research (on the mir-light induced anti-aging effects); Q.H., B.S. and C.S. designed the experiments, C.S. performed the experiments; B.S. and D.P. designed and performed the quantum chemistry calculations. All authors analyzed the results. and wrote the manuscript.

## Acknowledgments

This work was supported by the National Natural Science Foundation of China (T2394532, T2241002) and the National Key R&D Program of China (2021YFA1200404). The numerical computations were performed on the Hefei Advanced Computing Center.

## References

1 L. Partridge, J. Deelen, P. E. Slagboom, Facing up to the global challenges of ageing. Nature 561, 45–56 (2018).

2 B. K. Kennedy, et al., Geroscience: linking aging to chronic disease. Cell 159, 709–713 (2014).

3 C. López-Otín, M. A. Blasco, L. Partridge, M. Serrano, G. Kroemer, Hallmarks of aging: An expanding universe. Cell 186, 243–278 (2023).

4 M. S. Hipp, P. Kasturi, F. U. Hartl, The proteostasis network and its decline in ageing. Nat. Rev. Mol. Cell Biol. 20, 421–435 (2019).

5 T. B. L. Kirkwood, Understanding the odd science of aging. Cell 120, 437–447 (2005).

6 S. Ma, et al., Caloric restriction reprograms the single-cell transcriptional landscape of rattus norvegicus aging. Cell180, 984–1001 (2020).

7 A. R. Tari, et al., Neuroprotective mechanisms of exercise and the importance of fitness for healthy brain ageing. Lancet 405, 1093–1118 (2025).

8 D. Gems, L. Partridge, Stress-response hormesis and aging: “That which does not kill us makes us stronger”. Cell Metab. 7, 200–203 (2008).

9 D. E. Harrison, et al., Rapamycin fed late in life extends lifespan in genetically heterogeneous mice. Nature 460, 392–395 (2009).

10 M. Xu, et al., Senolytics improve physical function and increase lifespan in old age. Nat. Med. 24, 1246– 1256 (2018).

11 H. Zhang, et al., NAD+ repletion improves mitochondrial and stem cell function and enhances life span in mice. Science 352, 1436–1443 (2016).

12 A. Ocampo, et al., In vivo amelioration of age-associated hallmarks by partial reprogramming. Cell 167, 1719–1733 (2016).

13 V. N. Gladyshev, et al., Molecular damage in aging. Nat. Aging 1, 1096–1106 (2021).

14 A. A. Cohen, et al., A complex systems approach to aging biology. Nat. Aging 2, 580–591 (2022).

15 T. V. Pyrkov, et al., Longitudinal analysis of blood markers reveals progressive loss of resilience and predicts human lifespan limit. Nat. Commun. 12, 2765 (2021).

16 X. Liu, et al., Nonthermal and reversible control of neuronal signaling and behavior by midinfrared stimulation. Proc. Natl. Acad. Sci. U.S.A. 118, e2015685118 (2021).

17 J. Zhang, et al., Non-invasive, opsin-free mid-infrared modulation activates cortical neurons and accelerates associative learning. Nat. Commun. 12, 2730 (2021).

18 N. Li, et al., Demonstration of biophoton-driven DNA replication via gold nanoparticle-distance modulated yield oscillation. Nano Res. 14, 40–45 (2021).

19 Y. Yang, et al., AuNP-modulated qPCR: An optimized system for detecting MIR biophotons released in DNA replication. Chem. Eur. J. 29, e202203513 (2023).

20 C. Zhang, et al., Driving DNA origami assembly with a terahertz wave. Nano Lett. 22, 468–475 (2022).

21 C. D. De Magalhaes Filho, et al., Visible light reduces C. elegans longevity. Nat. Commun. 9, 927 (2018).

22 D. McCulloch, Body size, insulin/IGF signaling and aging in the nematode Caenorhabditis elegans. Exp. Gerontol.38, 129–136 (2003).

23 A. Picca, et al., Mitochondrial quality control mechanisms as molecular targets in cardiac ageing. Nat. Rev. Cardiol.15, 543–554 (2018).

24 S. N. Chaudhari, E. T. Kipreos, Increased mitochondrial fusion allows the survival of older animals in diverse C. elegans longevity pathways. Nat. Commun. 8, 182 (2017).

25 A. H. Khan, et al., Mitochondrial protein heterogeneity stems from the stochastic nature of co-translational protein targeting in cell senescence. Nat. Commun. 15, 8274 (2024).

26 L. C. Tábara, et al., MTFP1 controls mitochondrial fusion to regulate inner membrane quality control and maintain mtDNA levels. Cell 187, 3619–3637 (2024).

27 A. G. Vlassenko, M. M. Rundle, M. E. Raichle, M. A Mintun, Regulation of blood flow in activated human brain by cytosolic NADH/NAD+ ratio. Proc. Natl. Acad. Sci. U.S.A. 103, 1964–1969 (2006).

28 P. R. Kumar, et al., PGC-1α induced mitochondrial biogenesis in stromal cells underpins mitochondrial transfer to melanoma. Br. J. Cancer 127, 69–78 (2022).

29 J. Rodriguez, D. R. Larson, Transcription in living cells: Molecular mechanisms of bursting. Annu. Rev. Biochem. 89, 189–212 (2020).

30 A. E. Roux, et al., Individual cell types in C. elegans age differently and activate distinct cell-protective responses. Cell Rep. 42, 112902 (2023).

31 M. E. Papandreou, G. Konstantinidis, N. Tavernarakis, Nucleophagy delays aging and preserves germline immortality. Nat. Aging 3, 34–46 (2022).

32 T. K. Blackwell, et al., SKN-1/Nrf, stress responses, and aging in Caenorhabditis elegans. Free Radic. Biol. Med. 88, 290–301 (2015).

33 C. Y. Jao, A. Salic, Exploring RNA transcription and turnover in vivo by using click chemistry. Proc. Natl. Acad. Sci. U.S.A. 105, 15779–15784 (2008).

34 X. Xu, Y. Pang, X. Fan, Mitochondria in oxidative stress, inflammation and aging: from mechanisms to therapeutic advances. Signal Transduct. Target. Ther. 10, 190 (2025).

35 Y. Aman, et al., Autophagy in healthy aging and disease. Nat. Aging 1, 634–650 (2021).

36 M. P. Murphy, et al., Guidelines for measuring reactive oxygen species and oxidative damage in cells and in vivo. Nat. Metab. 4, 651–662 (2022).

37 E. Dehghan, et al., Hydralazine induces stress resistance and extends C. elegans lifespan by activating the NRF2/SKN-1 signalling pathway. Nat. Commun. 8, 2223 (2017).

38 A. Meléndez, et al., Autophagy genes are essential for dauer development and life-span extension in C. elegans. Science 301, 1387–1391 (2003).

39 J. T. Chang, et al., Spatiotemporal regulation of autophagy during Caenorhabditis elegans aging. eLife 6, e18459 (2017).

40 E. S. Kwon, S. D. Narasimhan, K. Yen, H. A. Tissenbaum, A new DAF-16 isoform regulates longevity. Nature 466, 498–502 (2010).

41 R. H. Houtkooper, et al., Mitonuclear protein imbalance as a conserved longevity mechanism. Nature 497, 451–457 (2013).

42 L. Yang, et al., Serine catabolism feeds NADH when respiration is impaired. Cell Metab. 31, 809–821 (2020).

43 F. Zhang, et al., Histone acetylation induced transformation of B-DNA to Z-DNA in cells probed through FT-IR spectroscopy. Anal. Chem. 88, 4179–4182 (2016).

44 S. Caine, et al., The application of Fourier transform infrared microspectroscopy for the study of diseased central nervous system tissue. Neuroimage 59, 3624–3640 (2012).

